# CRISPR-Cas immune repertoires as an ecological record of bacterial interactions with mobile genetic elements in the human gut

**DOI:** 10.64898/2026.03.18.712547

**Authors:** Ekaterina Avershina, Einar E. Birkeland, Cecilie Bucher-Johannessen, Trine Ballestad Rounge

## Abstract

Bacteria in the human gut influence host physiology and disease risk, but their ecology is strongly shaped by mobile genetic elements (MGEs) such as phages and plasmids. Past interactions between bacteria and MGEs can be inferred from CRISPR-Cas cassettes, which contain short DNA fragments derived from invading elements. To lay the groundwork for research on the impact of such interactions on the human host, we examined bacteria, MGEs, and CRISPR-Cas in the gut microbiome. Using fecal shotgun metagenomes from 1034 Norwegians, we constructed an extended microbiome resource comprising 1.7K prokaryotic mOTUs, 19.5K viral vOTUs and 24.2K plasmid PTUs. We also recovered 74.2K unique CRISPR-Cas cassettes to map past bacteria-MGE interactions and assessed their associations with human diet and lifestyle factors.

CRISPR-Cas spacers, and which viruses and plasmids they targeted, varied substantially within bacterial species, but were predominantly directed towards cohort-specific MGEs. Moreover, bacteria were more likely to target MGEs present in the same sample, consistent with local exposure. Bacteria also shared more targets within taxonomic families than across families, where mobilizable plasmids were more frequent among the targets. We did not find evidence that CRISPR-Cas spacers were related to characteristics of the human host, beyond the host-bacteria associations.

Together, this research provides a large-scale resource and a structured analysis of bacteria-MGE interactions in the gut microbiome, and their contribution to microbial ecosystem dynamics.

## Introduction

Human gut microbiomes are one of the most densely populated microbial environments, under intense ecological pressure and with frequent interactions between its inhabitants. To survive in such a competitive environment, bacteria must continuously adapt to selective pressures imposed by host immunity, nutrient availability, and other microbes. To respond to these fluctuating pressures, bacteria commonly exploit horizontal gene transfer, especially via plasmids and phages.^1,2^

At first glance, plasmids and phages seem to play opposite roles in the bacterial lifecycle, - plasmids often increase bacterial fitness, whereas phages exploit bacterial machinery for self-replication. Each, however, represent a mobile genetic element (MGEs) that carries both opportunity and risk for the bacterial host.^3,4^ On one hand, they drive adaptivity by disseminating genes involved in biofilm formation, virulence, toxin-antitoxin defense systems, and antibiotic resistance.^5–8^ On the other hand, they exert strong selective pressure through metabolic burden or cell death. To mitigate these antagonistic interactions, bacteria employ CRISPR-Cas immune system that targets invading and potentially deleterious DNA.^9,10^

The CRISPR-Cas immune system captures short DNA fragments from invading MGEs and retains them as spacers within the repeat-spacer-repeat arrays to rapidly recognize and fight off future invasions.^9^ These records can provide a window into the microecological context in which bacteria persist, including MGE populations they interact with.^11^ Variation in such contexts can translate into functional differences among closely related bacteria, including altered metabolic and physiological activity.

A substantial body of evidence links the composition and functional potential of the gut microbiome with human health outcomes, including gastrointestinal disorders, colorectal cancer, and a range of extraintestinal diseases.^12–15^ Although studies linking pathogenicity, CRISPR-Cas immunity, and MGEs do exist, direct mechanistic links between these processes and human health remain very limited.^16^

Despite the conceptual relevance of CRISPR-Cas systems for studying microbial interactions, they have barely been applied to investigate gut bacteria-virus-plasmid relationships in an integrated manner. We retrieved only three reports in PubMed that covered CRISPR-Cas interactions between gut bacteria and viruses^17–19^, and one that additionally touched upon plasmids^11^. Such paucity of studies reflects both methodological gaps and limited MGE databases, which hinder reliable spacer-target assignment and constrain inference of bacteria-MGE interactions.^20^ Consistent with this limitation, the largest database of CRISPR-Cas spacers available to date, SpacerDB, identified putative targets for only 10% of the spacers.^20^

To lay the groundwork for understanding the ecology of gut microbiome beyond its single constituents and its impact on human health, we characterize bacteria, viruses, plasmids, and CRIPSR-Cas cassettes from stools of 1034 adult individuals (Figure 1). We identify 140K novel CRISPR-Cas spacers, 17.4K novel plasmids, add bacterial host inference for 6K unknown-host or novel plasmids and provide these as a resource for the scientific community. We use this database to examine bacteria-MGE interactions by utilizing a historical record of CRISPR-Cas spacer repertoire, and to explore their associations to human host factors.

**Figure 1.**
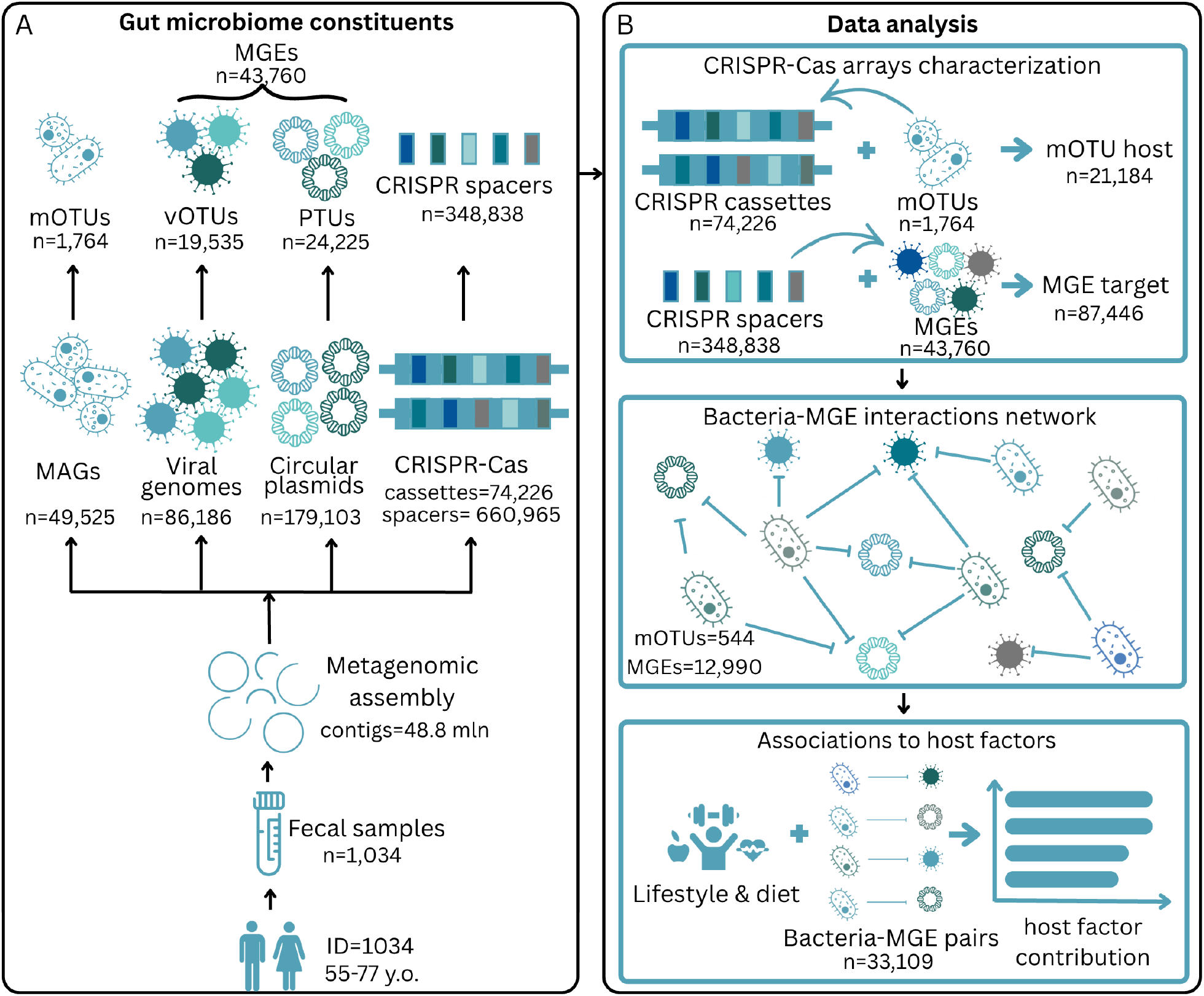
Overview of the study. A). Gut microbiome resource generation. The CRCbiome dataset comprises fecal samples from the Norwegian population (n=1034) aged 55-77 residing in South-East Norway. The samples were collected using a Fecal Immunochemical Test sampling kit. Shotgun metagenomic reads from these samples were assembled into 48.8 mln contigs. Prokaryotic metagenomic-assembled genomes and circular plasmids were generated from the contigs, and viral genomes and CRISPR-Cas cassettes were extracted. Prokaryotic genomes, viral genomes and circular plasmids were dereplicated to mOTUs, vOTUs and PTUs respectively based on sequence identity (mOTUs: 95%; vOTUs: 97.5%; PTUs: 90%). B). Data analysis. Bacteriome immunity towards vOTUs and PTUs was identified via CRISPR-Cas spacers’ target identification. The bacterial host was assigned if the cassette was located on a contig belonging to mOTUs. mOTU-vOTU-PTU networks were constructed, and MGE targets and mOTU-MGE interaction pairs were compared between individuals across different demographic and lifestyle factors. MGE: mobile genetic elements

## Materials and methods

All data analyses were performed using *pandas* v1.5.3, *numpy* v1.26.3, *statsmodels* v0.14.1, *scipy* v1.10.1 libraries, and visualized using *seaborn* v0.13.2 in Python v3.9.6 using Spyder IDE v5.3.0 unless stated otherwise.

### Study cohort

The dataset was previously described in^14,21^. The study used samples from the CRCbiome study, which took place between 2017 and 2022 and recruited participants from the FIT-arm of the Bowel Cancer Screening in Norway trial. All individuals were between the age of 55-71 and had tested positive for occult blood in their FIT sample (Eiken Chemicals Ltd., Tokyo, Japan) and were subsequently invited for colonoscopy. FIT samples were collected at home and shipped to the testing facilities (delivery time estimated 3-10 days). The leftover FIT-buffer was subjected to DNA extraction using the QIAsymphony automated system (Qiagen, Hilden, Germany) according to the manufacturer’s protocol with an additional off-board lysis procedure. Library preparation was performed with the Nextera DNA Flex Library Prep Reference Guide (Illumina, CA, USA) but scaling down to ¼ of the reference volume. Sequencing was performed on the Illumina NovaSeq X. In total, metagenomic profiles from 1034 individuals with a sequencing depth ≥1 Gbp were generated.

Additionally, participants filled in a Lifestyle and Demographic Questionnaire (LDQ) and a Food Frequency Questionnaire as described previously in Kværner et al.^22^ Relevant for this study, the LDQ contained questions regarding demographic factors (national affiliation, education, employment status, and marital status), and lifestyle factors (antacids use, smoking and snus habits, BMI, and physical activity level). The FFQ assessed habitual dietary habits over the preceding year. Based on the LDQ and FFQ, a healthy lifestyle index was calculated as described in Kværner et al.^23^ Prescription antibiotic use for each participant was obtained from the Norwegian Prescription Database (NorPD) and defined as having one or more prescriptions corresponding to any of the ATC codes J01, A07A, or P01A, four months prior to stool sampling.

### Metagenome assembled genomes dataset generation

Sequencing data was processed using metagenome ATLAS^24^ (v2.4.3) for assembly-based *de novo* generation of metagenome-assembled genomes (MAGs) and selection of representative genomes (mOTUs), as briefly described in^14^. Here, quality filtering was applied to paired-end sequencing reads using the BBMap^25^ (v37.99). Assembly was carried out using metaSpades^26^ (v3.13), using automatic k-mer size selection. Contigs were binned for the generation of MAGs using MetaBat^27^ (v2.2) and MaxBin^28^ (v2.14), using a minimum contig size of 1,500 and 1,000 bp, respectively. DAS Tool^29^ (v1.1) was used to resolve a non-redundant set of bins, using a score threshold of 0.5. Microbial community types for the samples were extracted from Birkeland et al^14^, where they were calculated using Dirichlet Multinomial Mixture models based on the bacterial relative abundance.

### Viral dataset generation

Viruses were recovered from ATLAS-generated contigs as described in Istvan et al.,^21^ with one notable modification. Unlike the previous publication, where only VirSorter2^30^ was used for viral sequence identification, here viruses were recovered using both VirSorter2^30^ (v2.2.2) and geNomad^31^ (v1.7.0). Only viruses identified by both were included in the final dataset. In case a viral sequence was identified within a longer contig, or if its location did not correspond between the two tools, the shorter sequence was assigned as viral. Viral genome sequences assigned a medium quality or higher (at least 50% genome completeness) as determined by CheckV^32^ (v0.8.1) were retained and dereplicated using Galah^33^ (v0.3.1), selecting the highest quality genome of the shortest possible length as the representative viral operational taxonomic unit (vOTUs). Taxonomy was determined using vConTACT2^34^ (v0.11.0) with the INPHARED database^35^ (version 01.05.2023). This analysis is a modified version of the VirMake pipeline^36^.

### Plasmid dataset generation

Circular plasmids were assembled with Sequence Contents-Aware Plasmid Peeler (SCAPP^37^ v 0.1.4) using minimal plasmid length of 1000 bp (default settings). Metagenome assembly graphs and BAM alignment files generated by read mapping to contigs were used as input. Assembled plasmids were then dereplicated into PTUs with *dedupe*.*sh* function from BBMap^38^ (v38.96) on 90%, 95% and 99% identity levels. After inspection, a 90% dereplication level was retained. Sequences matching the resulting PTUs were identified in the PLSDB database^39^ (version 10.02.2022) and the IMG/PR database^40^ (version 08.08.2023) using BLAST (v2.12). The results were then filtered, keeping only BLAST hits with ≥90% identity over at least 20% of the PTU length. Metagenomic reads from each sample were then mapped to a dereplicated plasmid database for relative abundance estimation. The plasmid was regarded as detected if metagenomic reads were mapped onto at least 75% of its length with a minimum of 90% identity. PTU mobility was derived using MOB-suite^41^ v3.1.9 typing, and antibiotic resistance genes were identified using RGI^42^ v6.0.5, allowing only strict and perfect hits. The workflow was organized into a pipeline using Snakemake^43^ v5.8.2.

### CRISPR-Cas dataset generation

CRISPR-Cas cassettes and their spacers were identified using CCTyper^44^ v1.8.0 with default settings in all metagenomic contigs and dereplicated PTU and vOTU datasets. CRISPR spacers were dereplicated at 99% identity level using cd-hit v4.8.1 and searched against local vOTU and PTU datasets, and against public databases of CRISPR spacers (CRISPR-Casdb^45^, version 06.04.2022, and SpacersDB^20^, version 10.05.2025), phages (INPHARED^35^, version 01.05.2023) and plasmids (PLSDB^39^, version 10.02.2022, and IMG/PR^40^, version 08.08.2023) using *blastn-short* in BLAST^46^ suite v2.12 with a minimum of 99% identity on at least 95% of the spacer’s length, to detect their potential targets. To avoid self-mapping, vOTUs and PTUs that contain CRISPR-Cas cassettes were excluded. The workflow was organized into a pipeline using Snakemake^43^ v5.8.2.

Rarefaction analysis for CRISPR-Cas spacer representativeness in cohort-specific and public databases was performed by randomly subsampling the databases with an increment of 1000 records until the size of the database was reached, 10 replicates per increment were generated. Individuality of spacers with regards to their targets was calculated as a median difference between the number of MGE targets present in an individual and the number of MGE targets present in ten other randomly selected individuals.

### mOTU-MGE interactions

In case CRISPR-Cas spacer with a known vOTU or PTU target was detected on a contig belonging to a known mOTU, the interaction between this mOTU and the target was recorded. All observed encounters of interactions between mOTUs and MGEs were then organized into a network. The observed mOTU-MGE network was visualized in Cytoscape^47^ v3.10.3. Pairwise mOTU connectedness was calculated based on presence of common CRISPR spacer targets between mOTUs.

To assess whether the observed bacteria-MGE interaction structure deviated from random expectations, we generated degree-preserving null networks using a bipartite configuration model. Weighted bacteria-MGE interaction counts were expanded into observation-level edges and randomized by double-edge swaps that preserved the total number of observations per mOTU and per MGE. The null model was initialized using 5 swaps per edge (burn-in phase) to ensure adequate randomization. The randomized edges were subsequently re-aggregated into weighted interaction pairs, and the number of unique bacteria-MGE pairs, interaction multiplicity distributions, and edge concentration metrics (Gini coefficient and Herfindahl-Hirschman index) were compared between the observed network and the null distributions using 200 permutations.

### Statistical analyses

Kruskal-Wallis test and ordinary least squares linear regression adjusted for sequencing depth were used for continuous data statistical analysis. Chi-square test and binomial distribution test were employed for categorical data analysis. Relative abundance correlations between mOTUs and MGEs were assessed using Spearman correlation. Only taxa with ≥20% prevalence in a domain-specific dataset were included. In case two taxa within a domain had ≥99% correlation to each other, only one of them was retained.

Comparative analysis and visualizations of Jaccard distances were performed in R (v4.2.0). Inter-individual similarity in MGE targeting was visualized using principal coordinates analysis (PCoA) and plotted with ggplot2 (v3.3.5). Difference in Jaccard distances was assessed using PERMANOVA implemented in the *adonis2* function of the vegan package (v2.6-2), using 999 permutations and the *by = “margin”* option to evaluate the independent contribution of each variable. The test was adjusted for the sequencing depth. In addition, a set of lifestyle, diet, and demographic variables’ contribution to mOTU sharing of MGEs was evaluated using individual PERMANOVA models. In this analysis, only individuals with at least one known mOTU-MGE pair were included (n=1031). Prior to analyses, missing values for host factors were imputed using the median for continuous variables and coded as missing for categorical values. Each model was adjusted for sequencing depth and microbial community type. The resulting R2 and p-values were summarized and visualized in barplots.

## Results

To assess bacterial immunity towards MGEs and its associations with human host factors, we constructed an extended microbiome resource of gut prokaryotes, viral and plasmid MGEs and CRISPR-Cas arrays from a CRC screening cohort^22^ comprised of 582 men and 452 women with a mean age of 67.0 ± 5.9 years. Prescription drug registry records showed that 141 individuals (13.6%) were prescribed antibiotics within 4 months prior to stool sampling. The metagenome dataset had an average sequencing depth of 3.3 ± 0.9 Gbp (12.0M ± 3.7M QC reads) per sample and was assembled into 48.8 million contigs with a mean length of 3,516 bp (SD=11,783 bp; min=500 bp; max=1.9 Mbp).

We have previously published a detailed description of read-based bacterial and archaeal composition of these samples in Birkeland *et al*.^14^, and of viral composition in Istvan *et al*.^21^, whereas plasmids and CRISPR-Cas were not previously reported. Here, we recovered 49,525 prokaryotic metagenome-assembled genomes dereplicated into 1,764 mOTUs (Bacteria, n=1,752; Archaea, n=12), and updated virus recovery using a combination of two tools for identification of viral sequences, resulting in 19,535 vOTUs, derived from 86,186 viral genomes. We also recovered 179,103 circular plasmids using SCAPP^37^ and dereplicated them into a set of 24,225 plasmid taxonomic units (PTUs). A detailed description of the prokaryome and virome can be found in Supplementary Text 1, and of the plasmidome in Supplementary Text 2. Additionally, pairwise diversity and relative abundance correlations between the microbiome constituents, as well as the overall microbial community composition assessments, are provided in the Supplementary Text 3.

### CRISPR-Cas cassettes exhibited high bacterial intraspecies variability

A total of 74,226 CRISPR-Cas cassettes were detected across all individuals (71±22 cassettes per sample), with the number of cassettes per sample largely determined by sequencing depth and assembly quality (Supplementary Figure 1A). Each CRISPR-Cas cassette comprised on average 12 repeats (SD=12; min=3; max=319) (Figure 2A), and the detection of the *cas* operon in addition to the repeat-spacer-repeat array depended on the contig length (Supplementary Figure 1B).

**Figure 2.**
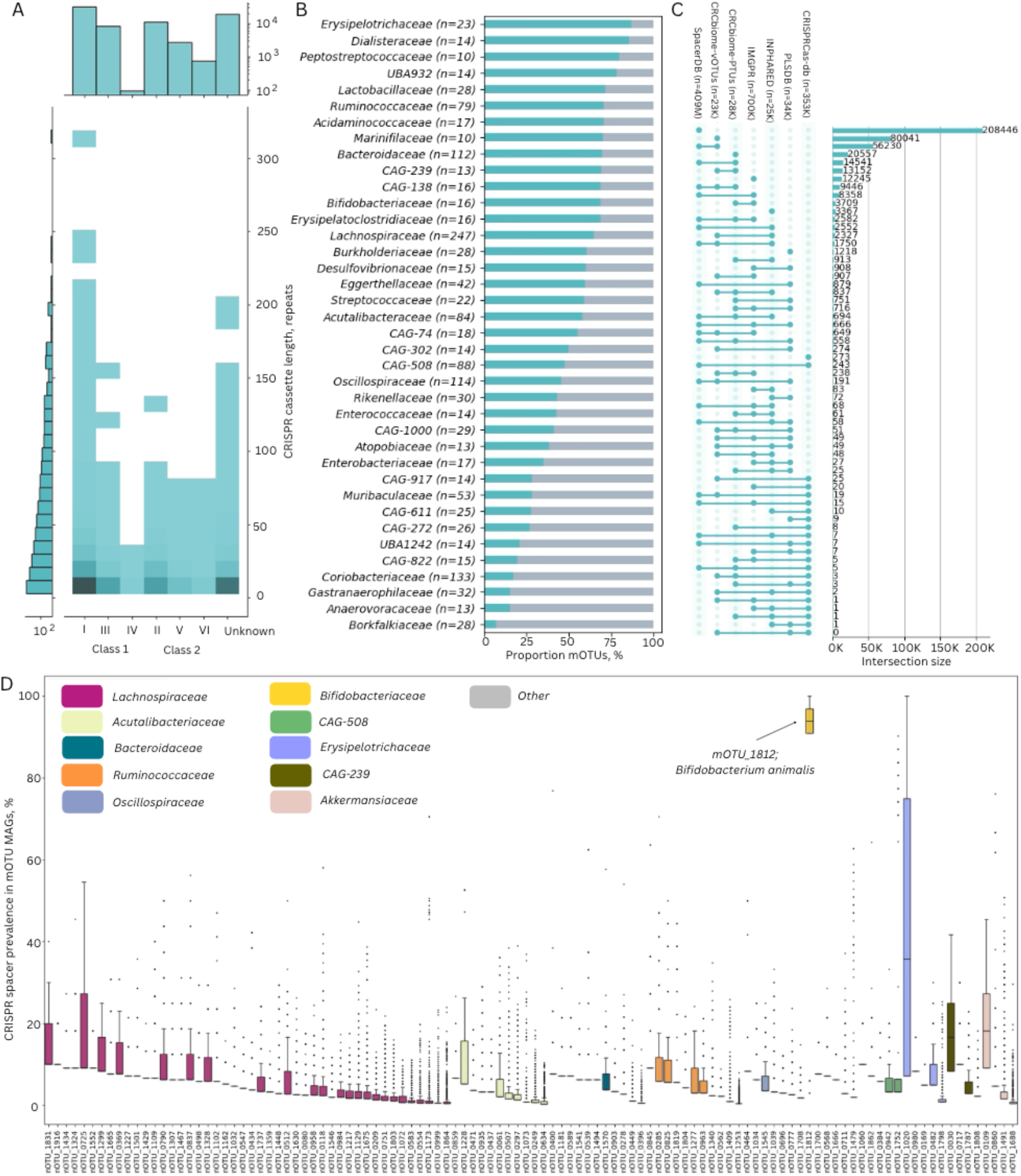
CRISPR-Cas dataset overview. A) Frequency of detection of CRISPR-Cas cassettes of each class and type, and number of repeats in these cassettes. B) Proportion of mOTUs belonging to a given taxonomy class where CRISPR-Cas cassettes were detected in at least one MAG (blue) or not detected in any MAG (grey). Only families with ≥10 mOTU representatives are depicted C) Number of CRISPR-Cas spacer clusters with ≥99% homology to records in public databases and to vOTUs and PTUs detected in the study. D) Heterogenity of CRISPR-Cas spacers in mOTUs. For each mOTU, the proportion of mOTU-MAGs that contained each CRISPR-Cas spacer detected in this mOTU, is depicted. All mOTUs with at least 10 MAGs, where CRISPR-Cas cassettes were detected, are included in the analysis.

Of the total identified CRISPR-Cas cassettes, 21,184 (28.5%) were located on contigs belonging to recovered genomes assigned to mOTUs. Overall, 916 mOTUs (51.9%) had at least one genome with an identified CRISPR-Cas cassette, and both the frequency of detection (Figure 2B) and the cassette length, i.e. number of CRISPR-Cas repeats (Supplementary Figure 1C), varied between bacterial families. There were 14 taxonomic families, with 1-3 mOTUs each, where no CRISPR cassettes were recovered (Supplementary Table 1). Since CRISPR-Cas cassettes have been reported to be encoded in plasmids and viruses^48,49^, we additionally searched for CRISPR-Cas cassettes in local PTU and vOTU sets. We found only 0.05% of PTUs (n=134) and 0.05% of vOTUs (n=109) to contain CRISPR-Cas cassettes.

For stringent identification of CRISPR spacers, we proceeded only with spacers from cassettes with a length of ≥10 repeats. This filtering excluded 91% of putative CRISPR-Cas cassettes and 24.3% of all identified spacers. Of the 660,965 remaining spacers, dereplication resulted in 348,838 unique CRISPR spacer clusters (average length 33±3 bp), 59.8% (n=208,446) of which were previously reported in the SpacerDB database (Figure 2C). Most dereplicated CRISPR spacers were singletons (n = 264,014, 75.7%), and most non-singleton spacers tended to be shared between individuals (n=81,792, 96.4%). Still, only 21 of those spacers were detected in >=10% population. Most non-singleton CRISPR spacers with known mOTU host (n=55,088) were detected in MAG bins belonging to the same mOTUs (n=51,535; 93.5%), and only 6.5% of them were detected in multiple mOTUs, mostly belonging to the same taxonomic family (Supplementary Figure 1D).

To assess the conservation of CRISPR-Cas cassettes within prokaryotes, we searched for shared spacers between MAGs within mOTUs with at least 10 CRISPR-Cas containing MAGs. Generally, mOTUs demonstrated high degree of heterogeneity in CRISPR spacers (Figure 2D). The one striking exception was mOTU1812 *Bifidobacterium animalis*, that shared most of its spacers in at least 30 of its 33 MAG genome bins. Such high conservation of spacers likely points towards a common origin of this strain. *B. animalis* is often added as a probiotic species in dairy products. Individuals with the *B. animalis* genome detected, had higher daily consumption of local fermented milk products which have *B. animalis* as a supplement (Supplementary Figure 2E, Kruskal-Wallis Hst=85.9, p=1.9*10^−20^), corroborating the recovery of a likely foodborne strain.

### CRISPR-Cas spacers targeted individual and population-specific MGEs, varying between microbial community types

To identify which MGEs CRISPR-Cas spacers targeted, we searched for homologous protospacers in public virus and plasmid databases, and the CRCbiome dataset-specific MGEs (Figures 2C, 3A).

**Figure 3.**
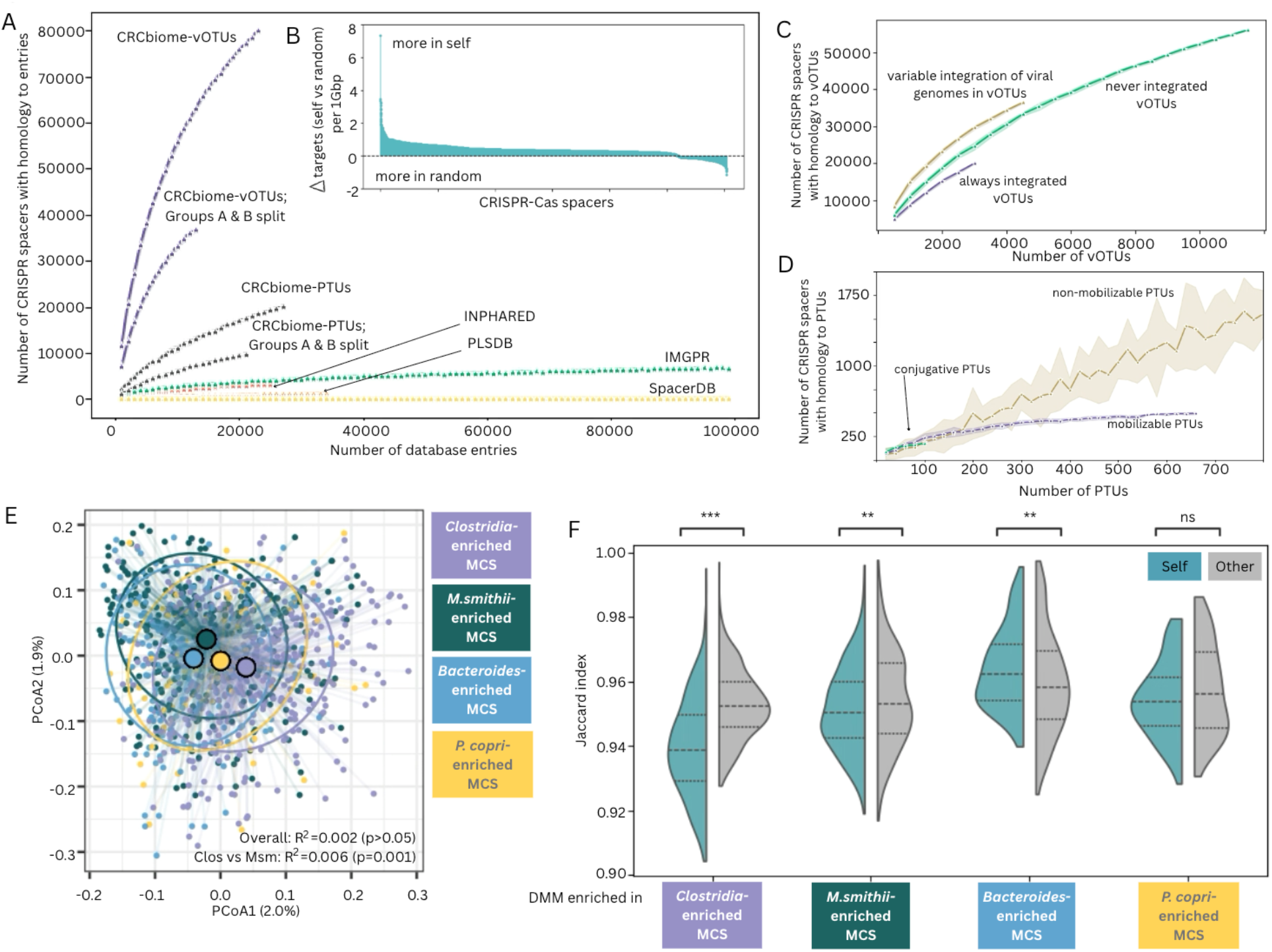
Targeting of MGEs by CRISPR-Cas spacers. A) CRISPR-Cas spacer clusters are better represented in the CRCbiome viral and plasmid datasets than in public viral, plasmid, and spacer databases. Rarefaction curves were calculated as the number of spacers with a homology to a sequence in a random subset of database sequences, using the mean of 10 randomizations and 95% confidence interval (shading), at increments of 1,000 database entries. To assess population specificity in MGE targeting, the CRCbiome individuals were randomly split in two equally sized groups, Groups A and B. Spacers detected in samples from Group A were then matched against the vOTUs and PTUs detected by read mapping in samples from Group B. The splitting was performed ten times for the calculation of the mean and of confidence intervals. B) Individuality of CRISPR-Cas targeting was assessed using the average difference in the number of MGE spacer targets detected in self vs other 10 randomly selected individuals. To account for sequencing depth, the number of detected targets in each sample was normalized to targets per 1 GBp sequencing data prior to difference calculation. For visualization purposes, only spacer clusters with targets detected either in self or in random individuals are included (n = 9,534, 10.9%); C) Enrichment of vOTUs by their host genome integration ability based on rarefaction analysis with 500 vOTUs increment. The average per 10 randomizations and 95% confidence interval (shading) are presented. D) C) Enrichment of PTUs by their mobility status based on rarefaction analysis with 25 PTUs increment. The average per 10 randomizations and 95% confidence interval (shading) are presented. (E) PCoA analysis of MGE targeting similarities between individuals contrasted by microbial community state defined in Birkeland et al.^14^ The similarity was assessed using Jaccard distance based on the fraction of spacers that shared common MGE targets. R^2^and p values are calculated using PERMANOVA analysis. MCS: microbial community state; Clos: Clostridia-enriched MCS; Msm: M.smithii-enriched MCS. (F) Difference in MGE targeting similarities between individuals within the same (blue) or different (grey) microbial community states defined in Birkeland et al^14^ The similarity was assessed using Jaccard distance based on the presence of spacers with common MGE targets, and varies from 0 (same targeted MGEs) to 1 (no overlap in targeted MGEs). MCS: microbial community state. ***p≤0.001; **p≤0.01; ns: not significant, p>0.05

A CRCbiome vOTU or PTU target was identified for 25.1% of CRISPR non-redundant spacers (n = 87,446), with a mean of 10 CRCbiome-MGE targets per spacer (SD=9, min=1, max=84). Spacers were more likely to target vOTUs or PTUs present in the same individual than those present in others (Kruskal-Wallis, p<1*10^−10^, Figure 3B), likely reflecting adaptation to an intra-individual gut environment. To assess whether there was an enrichment of targets in the CRCbiome dataset, we performed a rarefaction analysis excluding viruses and plasmids detected in the same individual from the matching dataset. MGEs present in other samples from the CRCbiome dataset were more commonly targeted than MGEs in unrelated public databases, potentially suggesting population specificity in addition to individual specificity (Figure 3A).

As with spacers, we also observed heterogeneity in MGE targeting at the mOTU level, assessing all mOTUs with at least 10 CRISPR-Cas containing MAGs. Each MGE was on average targeted by 9.8% of MAGs within each mOTU (SD=12.0%), and none were consistently targeted by all MAGs within any mOTU. At most, 41/47 (87.2%) CRISPR-containing genomes of the uncharacterized species UBA sp900544375 of the *Acutalibacteriaceae* family targeted the unknown virus vOTU-CB-12066. In addition, a set of viruses classified as *Akkermansia phage DTMo-2021a* were targeted by spacers detected in ~85% of *Akkermansia* sp001580195 and *Akkermansia muciniphila* B genomes.

Overall, vOTUs where at least one, but not all, viral genomes were integrated into bacterial host genomes, were more frequently targeted than those vOTUs where viral genomes were either never or always integrated (Figure 3C). Similarly, PTUs with no genes enabling plasmid mobility or conjugation (non-mobilizable PTUs) were more frequently encountered among CRISPR spacer targets than those that had such genes (Figure 3D).

CRISPR-Cas targeting of MGEs reflects prior exposure to such MGEs, and the collective record of targeted elements within a sample could be seen as a reflection of long-term exposure. We hypothesized that the collective record of MGE exposure encoded by CRISPR spacers would be determined by which bacteria harbor them. We therefore evaluated to what extent microbial community states derived from prokaryote abundances (defined in Birkeland et al.^14^) were associated with differences in the overall pattern of MGE targeting in a sample. Although overall MGE targeting by CRISPR-Cas did not significantly reflect the community states, individuals enriched in either *Clostridia* or *Methanobrevibacter smithii*, resembled other members of the same group more than individuals from other community states (Figures 3E & 3F). Interestingly, individuals with the *Bacteroides* spp.-enriched microbial community state showed an opposite pattern, potentially reflecting higher diversity among the dominant bacterial species.

### Bacteria tended to share CRISPR-Cas MGE targets with other bacteria belonging to the same taxonomic families

Both plasmids and phages show non-random bacteria host association, and host range spans a continuum from highly restricted strain-specific phages to relatively generalist broad-host-range plasmids or polyvalent phages.^50–52^ CRISPR-mediated targeting of MGEs provides evidence of prior infections or contact between bacteria and MGE, and we therefore set out to elucidate mOTU-MGE networks based on these patterns of recognition.

There were 146,256 non-redundant CRISPR-Cas spacers with at least one representative located on contigs belonging to recovered mOTUs. Out of these, 35,887 (24.5%) had targets identified among CRCbiome vOTUs and/or PTUs (excluding those with CRISPR-Cas systems). Overall, we recovered 27,062 unique mOTU-vOTU and 6,047 mOTU-PTU target pairs with 544 mOTUs, 10,519 vOTUs and 2,471 PTUs. This observed number of mOTU-MGE pairs is significantly lower than what would have been expected under the pretense of random pairing of these constituents (expected=116,838±103, p=0.005). Moreover, most of the observed mOTU-MGE pairs were encountered more often than expected (mean multiplicity observed = 4.7±11.7 vs expected = 1.3±0.01; p=0.005), and with a high degree of inequality between the pairs’ detection (Gini inequality coefficient observed = 0.65 vs expected = 0.22±0.00; p=0.005), strongly supporting selectivity in bacterial immunity towards specific MGEs.

Interactions defined by mOTU-MGE pairs tended to form subnetworks comprising taxonomically related mOTUs sharing multiple vOTUs/PTUs (Figure 4). However, some bacteria, such as mOTUs belonging to *Acutalibacteriaceae*, had similar connectivity to mOTUs from other families (Supplementary Figure 2). vOTUs and PTUs were characterized using their integrative status and mobility traits. Viruses that were found integrated in the host genome were targeted by a narrower range of bacteria than those without evidence of integration (χ^2^=73.7; p<1*10^−10^). For plasmids, those that were mobilizable were targeted by a broader range of bacteria compared to non-mobilizable or conjugative ones (χ^2^=84.6; p<1*10^−10^).

**Figure 4.**
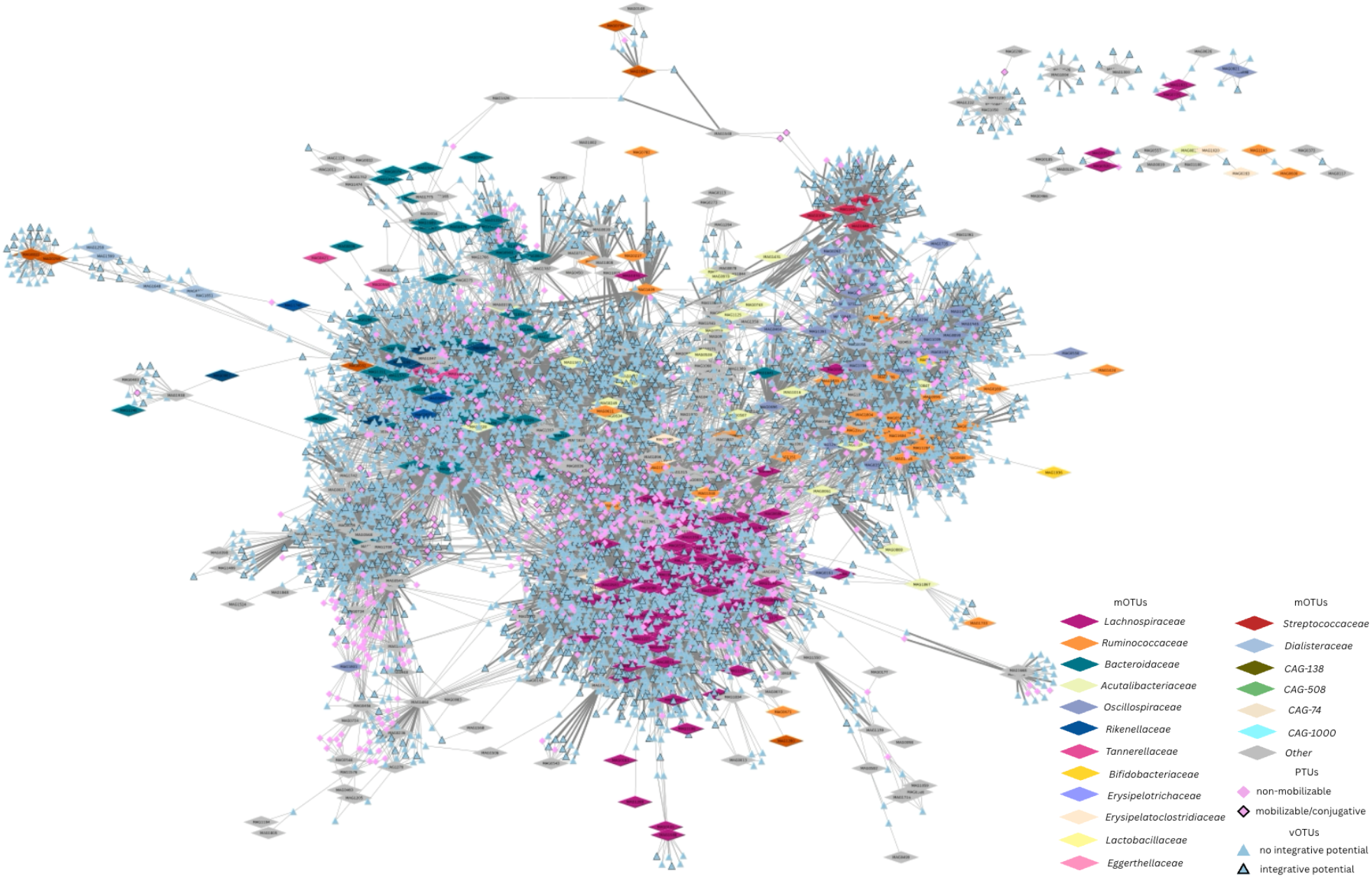
mOTU-MGE interaction network. Each diamond represents an mOTU colored by a taxonomic family. Blue triangle codes vOTUs, pink diamonds - PTUs. Each line represents a connection via a spacer, with the thickness depicting whether the connection was detected in ≥5 individuals (thick line) or <5 individuals (thin line). For visualization purposes, only interactions that connect at least 2 mOTUs, are depicted.

### Prokaryotic immunity mirrors associations between microbial constituents and host factors

CRISPR-Cas spacers provide a record of past MGE invasions, thus extending the information obtained from a single microbiome snapshot to include recent history. In fact, the targets of most CRISPR-Cas spacers with a known MGE target were below the detection limit (n=78,482, 89.7%). Incorporating this signal could reveal gut microbiome-host factor associations that are otherwise overlooked. Furthermore, the presence of a shared target within the same mOTU may reflect similarities in the microenvironments occupied by its members. For two bacteria within the same mOTU to share a target, both must have been exposed to the same invading MGE, implying physical and ecological proximity.

To evaluate the environmental influence on mOTU sharing of MGE targets across the gut microbiome, we constructed a distance matrix by performing a pairwise comparison of the fraction of mOTU-MGE target pairs that were shared for each pair of participants. We tested whether mOTU-specific sharing of targets was associated with host demographic, lifestyle, or dietary variables and compared these patterns to associations with beta-diversity based on the presence or absence of mOTUs, vOTUs, and pOTUs. mOTU-MGE targeting-derived associations seemed to mirror individual microbial constituents’ associations, especially that of prokaryotes, also when adjusting for the microbiome community state and sequencing depth (Figure 5). In line with this, no statistically significant associations were observed for any commonly targeted MGEs (MGEs targeted by at least 50 genomes within an mOTU, Supplementary Table 2), which suggests that prokaryote immunity *per se* is not influenced to a significant extent by lifestyle factors.

**Figure 5.**
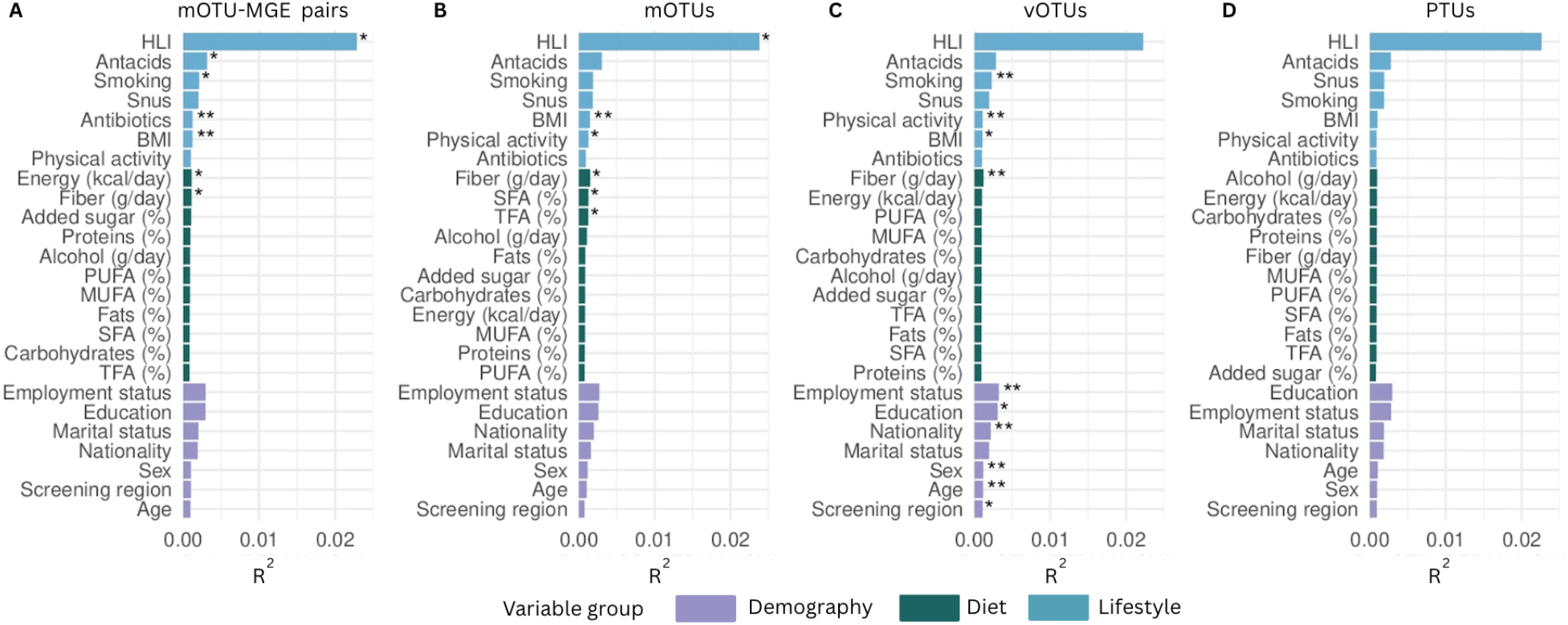
Associations between individuals’ microbial community similarities and human host factors. Similarities were assessed using Jaccard index calculated based on the presence/absence of (A) mOTU-MGE pairs (presence/absence of genomes within the same mOTUs with a CRISPR-Cas immunity towards the same MGEs); (B) mOTUs; (C) vOTUs and (D) PTUs. *PERMANOVA p<0.05; **PERMANOVA p<0.01. PERMANOVA analysis was adjusted for the sequencing depth and microbial community states. HLI: Healthy Lifestyle Index; BMI: Body Mass Index; SFA: Short-chain fatty acids; TFA: Trans Fatty Acids; MUFA: Monounsaturated fatty acids; PUFA: Polyunsaturated fatty acids; (%): Percentage of total energy intake per day.

## Discussion

In this work, we identified and characterized the prokaryome, virome, plasmidome, and CRISPR-Cas repertoires from 1,034 shotgun metagenomes, and present this as a comprehensive microbiome resource for the community to enable comparative analyses, identification of novel taxa and functions, and exploration of community-level and cross-entity interactions. Using this resource, we characterized bacteria-MGE interactions at a population and individual scale, and revealed a highly individualized landscape of bacterial immunity, which appears to reflect microbial ecology within the gut rather than host-associated factors.

Despite the rapid growth of public metagenomic sequencing, a substantial fraction of gut microbial diversity remains uncharacterized.^53^ This “dark matter” is particularly pronounced for MGEs, where many sequences lack confident taxonomy and host annotation. Large-scale genome- and virome-catalog efforts continue to expand available reference space, yet they consistently report extensive novelty and limited host resolution for many recovered viral and plasmid sequences.^40,54^ By leveraging CRISPR-Cas spacers as records of prior encounters between bacteria and MGEs alongside bacterial, viral and plasmid genomes, we provide empirical links between bacterial hosts and their associated viruses and plasmids, thereby offering host-range hypotheses at the species-, genus-, and family-level.

For a CRISPR-Cas spacer to be incorporated into a CRISPR array, several sequential events must occur: (i) an MGE must invade the bacterial cell, (ii) the CRISPR adaptation machinery must be triggered, and (iii) a segment of the invader’s DNA must be sequentially integrated as a new spacer into the host CRISPR locus, where multiple regions of an invading genome can serve as potential protospacer templates.^55,56^ As a consequence, each individual’s CRISPR spacer repertoire reflects not only bacteriome variation but also personal history of exposures to distinct viral and plasmid MGEs.^57,58^ Consistent with this, we observed marked interindividual variation in CRISPR spacer content, with most spacers being detected in only one or a few individuals. This high degree of personalization is likely a result of bacterial adaptation to local MGE exposure and could serve as a fingerprint of microevolutionary dynamics within the gut. A notable deviation was the relatively conserved spacer repertoires in *Bifidobacterium animalis* genomes recovered from 33 individuals. These individuals reported higher consumption of dairy products typically containing *B. animalis*, consistent with diet-driven species acquisition.^59,60^ Although spacer variability within this species was markedly lower than that observed for other taxa, likely reflecting a common lineage, detectable differences in CRISPR content were still present. This indicates that even shared, diet-associated strains undergo individual-specific diversification following colonization. Apart from this observation, we did not observe other host lifestyle-related patterns in MGE targeting by bacteria.

The pronounced bacterial adaptivity to local MGE exposure was further supported by the higher detection rate of MGE targets in the cohort-specific database compared to public repositories. Cross-kingdom network analyses are typically inferred from co-occurrence patterns derived from relative abundances, reflecting predominantly indirect or environmentally driven associations rather than biological interactions.^61^ In contrast, CRISPR-Cas spacers provide records of prior encounters between bacteria and MGE, and enable the network reconstruction grounded in direct molecular evidence. We found that bacteria exhibited strong selectivity in incorporating CRISPR-Cas spacers targeting specific MGEs, with bacteria within the same taxonomic families tending to share overlapping MGE pools.

By contrast, mobilizable plasmids and non-integrated viruses, exhibited interactions indicative of a broader bacterial host range. These genetic elements are intrinsically more ‘selfish’ and likely selected for maximizing horizontal transmission across diverse hosts, rather than long-term alignment with any single host lineage, whereas prophages, and possibly non-mobilizable plasmids rely more strongly on their hosts’ fitness.^62,63^ Consistent with this, CRISPR-Cas targeting varied by MGE lifestyle and mobility. Phages with intermittent integration into bacterial genomes were more frequently targeted than those that were always integrated or never integrated. This observation aligns with the report that bacteria more readily acquire spacers against viruses entering a lysogenic state upon infection, as a purely lytic phage infection may leave insufficient time for spacer acquisition, while targeting a stable prophage carries the risk of autoimmunity.^64^ Similarly, non-mobilizable plasmids showed higher targeting rates than mobilizable or conjugative ones.

Our study has several strengths. In addition to standard metagenomic assembly, we systematically recovered viral genomes and independently assembled bacterial plasmids, enabling a high-resolution characterization of MGEs. We further integrated these with CRISPR-Cas spacer profiles to investigate bacterial adaptation to individual MGEs. Lastly, the inclusion of participants with extensively validated dietary, lifestyle, and demographic data allowed us to explore associations between CRISPR-Cas immunity and host-related factors. Several limitations should be acknowledged. First, FIT samples contain ~10 mg of stool. While suitable for microbiome profiling, the limited input material, combined with the modest sequencing depth, has likely reduced the recovery of genomes from low-abundance taxa.^21,65,66^ Second, we did not account for spacer chronology to distinguish between recent and more ancient bacteria-MGE interaction events. Finally, our analysis focused on viruses and plasmids and did not include other classes of MGEs that may also be targeted by CRISPR-Cas systems.

## Conclusion

Integrating prokaryotic, viral, plasmid, and CRISPR-Cas data from >1000 individuals, we provide a comprehensive population and individual-scale resource for investigating MGEs and their interactions in the gut. This includes spacer-based bacterial host ranges for isolated MGEs. Our analyses reveal a highly individualized and cohort-specific landscape of CRISPR-based immunity that reflects bacterial host microenvironment more strongly than human host characteristics. CRISPR spacer-informed network reconstruction uncovered selective MGE targeting and shared MGE pools within bacterial families that would not be detectable through co-occurrence networks. Variation in host range based on MGE lifestyle and mobility further suggests evolutionary trade-offs between MGEs’ horizontal dissemination and longterm host alignment. This work provides a valuable foundation for future investigations linking microbial evolutionary dynamics and ecosystem stability.

## Data availability

DNA sequencing data generated in this study are deposited in Federated EGA under accession code EGAS50000000170. Per participant consent, submitted FASTQ files exclude reads mapping to the human genome. Processing of data from this study must comply with the General Data Protection Regulation (GDPR). Access can be obtained by following the procedure described here: https://www.mn.uio.no/bils/english/groups/rounge-group/crcbiome/.

DNA sequences of the GM constituents detected in at least 5 individuals, and all related metadata are provided in FigShare (doi:10.6084/m9.figshare.31707688), and scripts are available at https://github.com/Rounge-lab/CRISPR_interactions. The associated metadata include genome length, completeness, and the number of genomes used to construct each taxonomic unit. For viruses, we report detected auxiliary metabolic genes, whereas for plasmids we provide information on mobility, antibiotic resistance genes, and predicted bacterial hosts based on reference databases or relative abundance patterns. We indicate whether CRISPR-Cas systems were identified in each mOTU. For MGEs, we report whether they were targeted by recovered CRISPR-Cas spacers, whether they encoded CRISPR-Cas systems themselves, and their spacer-inferred potential hosts. Additionally, we provide all detected mOTU-MGE target pairs along with their frequency of occurrence.

## Author contributions

EA, EB, TBR - conceptualization; EB, CBJ - data generation; EA, EB, CBJ - data curation, formal analysis & visualization; EA - original draft writing; EA, EB, CBJ, TBR - draft review & editing; TBR- funding acquisition; TBR - supervision & project administration

## Acknowledgements

We would like to express our gratitude to Jan Inge Nordby from the Department of Medical Biochemistry at Oslo University Hospital, Erik Natvig and Anita Jørgensen from the Department of Colorectal Cancer Screening at the Cancer Registry of Norway (CRN), National Institute of Public Health (NIPH), and Vahid Bemanian from the Pathology Department at the Akershus University Hospital for their contributions to biobanking, laboratory work, and data management. We also wish to thank Harri Kangas and Pekka Ellonen at the Finnish Institute of Molecular Medicine (FIMM) for their assistance in preparing sequencing data; library preparation and sequencing were conducted at the FIMM Technology Centre, supported by HiLIFE and Biocenter Finland. Additionally, we acknowledge the support of current and former group members for their administrative and scientific contributions, including Maja Sigerseth Jacobsen, Paula Istvan and Paula Berstad. The CRCbiome study used in this publication, was supported by the Norwegian Cancer Society, projects 190179 (TBR), 297310 (TBR), and 198048 (Paula Berstad, Cancer Registry of Norway, Norwegian Institute of Public Health, Oslo, Norway), and the South-East Norway Regional Health Authority projects 2022067 (TBR) and 2020056 (TBR).

In this work, ChatGPT LLM was used for assistance with scripting, and to improve language and readability. All AI-assisted text revisions and code suggestions were reviewed, evaluated and finalized by the authors. Figure panels were assembled in *Canva*, and explanatory text and annotations were added without altering the original graphs.

## Abbreviations

CRISPR-Cas: Clustered Regularly Interspaced Palindromic Repeats (CRISPR) and CRISPR-associated proteins (Cas)
IMG/PR: Plasmid database derived from the Integrated Microbial Genomes & Microbiomes
IQR: interquartile range
MAG: Metagenome-Assembled Genome
MCS: microbial community state
MGE: Mobile Genetic Elements
mOTU: metagenomic Operational Taxonomic Unit
PCoA: Principal Coordinate Analysis
PLSDB: The Plasmid Database by NCBI
PTU: Plasmid Taxonomic Unit
SD: standard deviation
vOTU: viral Operational Taxonomic Unit

