## Supplementarl Info for "CRISPR-Cas immune repertoires as an ecological record of bacterial interactions with mobile genetic elements in the human gut"

**Supplementary Text 1. Description of bacteriome/archaeome and virome data**

**Supplementary Text 2. Description of plasmidome data and its correlations with prokaryotes and ARGs**

**Supplementary Text 3. Extended microbiome diversity and co-occurrence**

**Supplementary Figures**

*Supplementary Figure 1. CRISPR-Cas dataset characteristics*

*Supplementary Figure 2. Connectedness between mOTUs*

*Supplementary Figure 3. Bacteriome/archaeome and virome overview*

*Supplementary Figure 4. Technical characteristics of the plasmidome data*

*Supplementary Figure 5. Plasmidome overview*

*Supplementary Figure 6. Extended microbiome characteristics*

**Supplementary Tables**

*Supplementary Table 1. Proportion of mOTUs with CRISPR-Cas cassettes detected*

*Supplementary Table 2. Differential presence of MGE targeting in mOTUs based on host demographics and lifestyle factors*

**Supplementary Text 1. Description of bacteriome/archaeome and virome data**

The dataset comprised 1764 mOTUs (Bacteria, n=1752; Archaea, n=12) and 19 535 vOTUs, derived from 49 525 prokaryotic MAGs and 86 186 viral genomes, respectively. *Clostridia* was the most common class of recovered mOTUs (n=738), and along with *Bacteroidia* (n=245), these included the most prevalent prokaryotic taxa (Supplementary Figure 3A). Average genome completeness of mOTUs ranged from 84.5% ± 14% (class *Thermoplasmata*) to 97.1% ± 5.4% (class *Fusobacteriia*) and average genome length varied from 1.4 Mbp ± 0.3 Mbp (class *Thermoplasmata*) to 4.4 Mbp ± 0.6 Mbp (class *Lentisphaeria*). Mean completeness of vOTUs ranged between 70.9% ± 14.1% (*Winoviridae*) and 95.4% ± 9.8% (*Inoviridae*), and their length varied from 5.2 Kbp ± 1.5 Kbp (*Microviridae*) to 119 Kbp ± 40.7 Kbp (*Steigviridae*) (Supplementary Figure 3B). Only 5.6% of vOTUs (n=1102) were assigned taxonomy on a family level, and *Microviridae* comprised the most vOTUs (n=477). Viral genomes of 3131 vOTUs (16.0%) were always classified as integrated into bacterial genomes, 4753 vOTUs (24.4%) had at least one but not all viral genomes integrated, and 11651 vOTUs (59.6%) had no genomes integrated into bacterial genomes.

**Supplementary Text 2. Description of plasmidome data and its correlations with prokaryotes and ARGs**

In total, 179,103 plasmids (86±32 per sample, min=20, max=336) were recovered by SCAPP[30]. The number of recovered plasmids was influenced by sequencing depth, assembly quality, and prokaryotic richness (Spearman rho=0.28-0.45; Supplementary Figure 4A). We performed intra-dataset dereplication of plasmids to define a dataset-specific non-redundant catalogue. Regardless of the similarity threshold cutoff (90%, 95% or 99%), dereplication resulted in a set of 24,225 non-redundant plasmid taxonomic units (PTUs), indicating high genomic stratification. The boundary between plasmids and phages is somewhat blurred due to the existence of phage-plasmids, - mobile genetic elements that exhibit features of both. We assessed the extent to which our plasmids are also classified as phages using VirSorter2. We found 22.8% of dereplicated plasmids (n=5,521) to also be classified as phages.

Comparing our PTUs with existing databases, only 6 % of the PTUs (n=1360) matched a reference sequence in the PLSDB database, which is largely composed of plasmids derived from sequencing individual bacterial genomes deposited in the NCBI. The IMG/PR database additionally comprises sequences of plasmids detected in metagenomes (constituting 78.9% of the database). Here, 27.9% of PTUs (n=6,779) matched IMG/PR plasmids, where 89.6% of these matching IMG/PR plasmids (n=6,080) were metagenome-derived. Almost 60% of the IMG/PR plasmids that PTUs had homology to, were associated with the human digestive system according to IMG/PR metadata (n=3,483), and their representation in the dataset was significantly more frequent than generally in the IMG/PR database (binomial $p<1*10^{-10}$; Supplementary Figure 5A). PTUs had an average length of 6,753 bp ± 18,992 bp (min: 1,000 bp; max: 507,676 bp) and tended to either be shorter or of the same length as their closest match IMG/PR reference plasmids (Supplementary Figure 5B), although there was a minor fraction of PTUs that were at least 2 times longer than their respective references (n=58; 0.2%), possibly reflecting challenges arising with circular plasmid assemblies from short reads (Supplementary Figure 4B).

4490 PTUs matched IMG/PR reference plasmids with known bacterial host (66.2% of all PTUs with a homology to IMG/PR plasmids), including 3442 PTUs (50.8%) with known host family delineation. We set to infer bacterial host taxonomy for previously unidentified plasmids taking advantage of the known bacterial composition of the dataset by defining the best-correlated mOTU/PTU pairs based on their relative abundances in the samples. To assess this method, we used PTUs with known host family prediction as provided by IMG/PR. For each such PTU, we selected the best-matching mOTU based on the highest Spearman correlation coefficient $rho_{max}$, and assessed the family correspondence between IMG/PR prediction and mOTU family delineation (Supplementary Figure 5D). PTU/mOTU pairs with mOTU family corresponding to IMG/PR host prediction tended to have higher correlation coefficient with an average $rho_{max}$=0.4 (SD=0.2) compared to those pairs where taxonomy between best-matching mOTU and IMG/PR host prediction did not correspond ($rho_{max}$=0.29; SD=0.21) (Kruskal Wallis Hst=116.8, $p<1*10^{-10}$; Supplementary Figure 4C). The precision of such host taxonomy inference ranged between 73% ($rho_{max}$≥0.1) and 84% ($rho_{max}$≥0.3) (Supplementary Figure 4D). Using $rho_{max}$≥0.3, we additionally inferred host taxonomy to novel PTUs that lacked any IMG/PR reference plasmids and to PTUs that matched a plasmid with no host taxonomy reported (n=6,318). Majority of these PTUs were predicted to be hosted by *Lachnospiraceae* (n=2,023, 32.0%), *Bacteroidaceae* (n=596, 9.4%) and *Oscillospiraceae* (n=594, 9.4%).

When assessing the potential for horizontal gene transfer abilities, only a minor fraction of PTUs were mobilizable (2.8%, n=679) or conjugative (0.4%, n=105), with the rest considered to be non-mobilizable. *Enterobacteriaceae*, *Lactobacillaceae* and *Desulfovibrionaceae* had the highest proportion of mobilizable and conjugative PTUs (Supplementary Figure 5B). Conjugative PTUs tended to have a higher proportion of ARG-encoding PTUs ($\chi^2$=240.2; $p<1*10^{-10}$) with a higher number of ARGs ($p_{adj}$=0.001) and efflux pumps ($p_{adj}$=0.02) conferring resistance to more drug classes ($p_{adj}=5.8*10^{-5}$) (Supplementary Figure 5C). Overall, PTUs comprised ARGs conferring resistance to drugs across 12 antibiotic classes, including beta-lactams (3rd gen cephalosporins), glycopeptides (vancomycin), macrolides (erythromycin), and aminoglycosides (gentamycin).

**Supplementary Text 3. Extended microbiome diversity and co-occurrence**

Based on read mapping, we detected a mean of 186±51 mOTUs, 112±38 vOTUs and 1147±360 PTUs per individual, with strong correlation in alpha diversity between the constituents (Supplementary Figure 6A). Individuals with recent antibiotics use (within 4 months prior to sampling), had lower diversity across microbial constituents (Supplementary Figure 6B). Moreover, it was increasing by around 3 mOTUs, 1 vOTU and 16 PTUs with each year passing since the last antibiotic use (Supplementary Figure 6C). The diversity of PTUs with ARG load, however, seemed to be independent of time since the last use of antibiotics ($p_{adj}$>0.05).

When combining all microbiome constituents, significant differences in composition were detected with regards to recent antibiotic use (within 4 months prior to sampling; Supplementary Figure 6D) (Jaccard and Bray-Curtis respectively, PERMANOVA p<0.01).

Spearman correlation between mOTUs and vOTUs ranged between rho=-0.29 (vOTU-CB-07736, Unclassified to mOTU0825, *Ruthenibacterium lactatiformans*) and rho=0.91 (vOTU-CB-03962, Unclassified to mOTU0735, *Alistipes putredinis*). Most mOTU-vOTU pairs were weakly correlated based on their relative abundances (median rho=0.001, 25th percentile rho=-0.02, 75th percentile rho=0.03). Among 38 vOTUs with strong Spearman correlation to mOTUs (|rho|>0.5), 30 vOTUs had viral genomes integrated into contigs within MAGs (Supplementary Figure 6E).

Positive Spearman correlation between relative abundances of mOTUs and PTUs was used for PTU host assignment and is provided in Supplementary Text 2. Most PTUs with negatively correlated relative abundance to that of mOTUs, had rather weak correlation (median rho=-0.03, IQR=0.03). Among PTUs with rho<-0.1, majority of PTUs with known family delineation (n=5790) were predicted to be hosted by *Lachnospiraceae* mOTUs (8.4%, n=488). Almost 80% of these *Lachnospiraceae* PTUs (n=388) had a negative correlation to mOTUs within several taxonomic families, mostly within *Lachnospiraceae* (n=381 PTUs), *Oscillospiraceae* (n=325 PTUs) and *Ruminococcaceae* (n=286 PTUs).

### Supplementary Figures

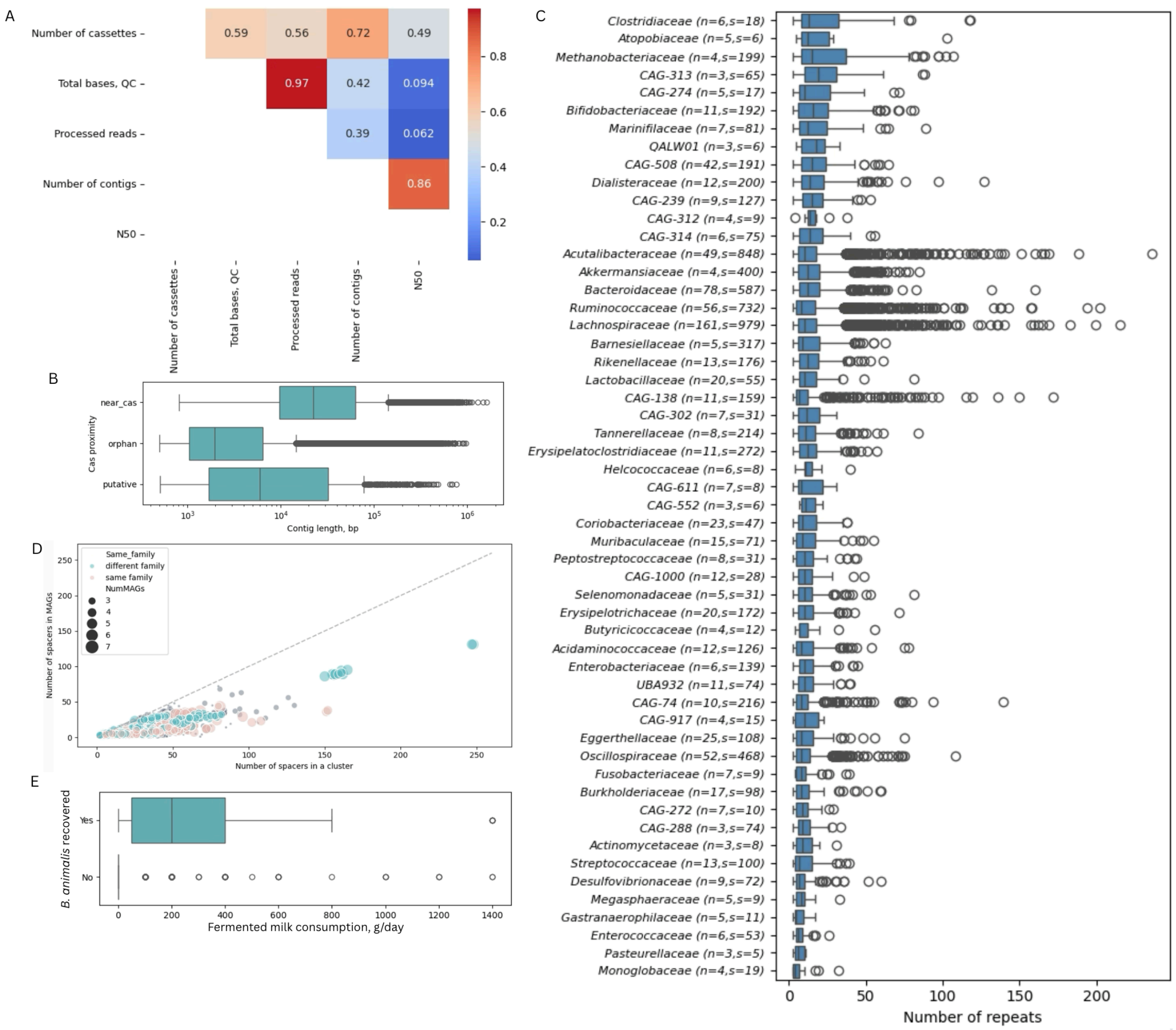


*Supplementary Figure 1. CRISPR-Cas dataset characteristics A) Pearson's correlation between number of CRISPR-Cas cassettes detected per sample vs sequencing depth, number of contigs and assembly quality (N50). B) Length of metagenomic assembly contigs with CRISPR-Cas cassettes detected near cas genes, separate from cas genes (orphan), or putative cassettes. C) CRISPR-Cas cassettes length (number of repeats) of cassettes stratified by MAGs taxonomic family. Only bacterial families where CRISPR cassettes were detected in at least 3 mOTUs, and in samples from at least 5 individuals, were included. D) Number of CRISPR-Cas spacers that were located on MAGs within mOTUs vs the full CRISPR-C as spacer cluster size (number of spacers in the dereplicated cluster). Size depicts number of different mOTUs that the recovered MAGs belonged to E) Daily consumption of fermented milk products by individuals from which B. animalis genome was recovered (n=33) and those where B. animalis was not recovered (n=1001).*

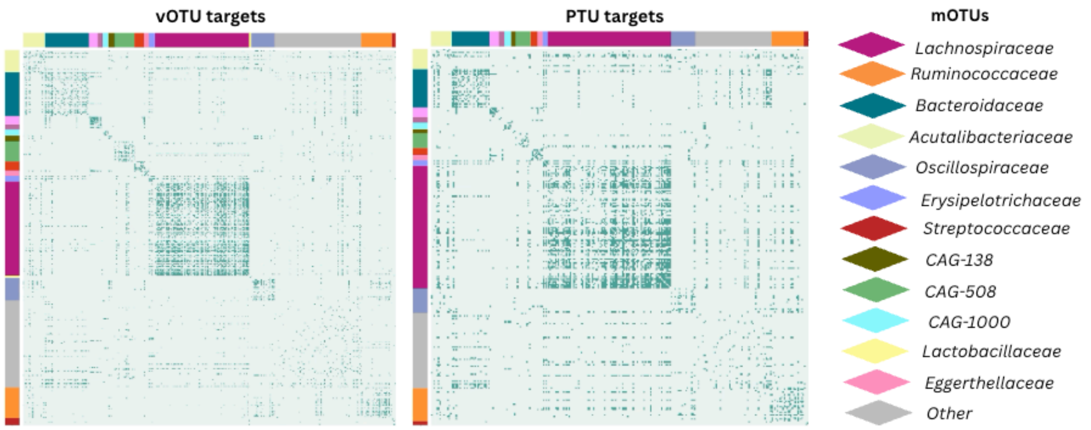


*Supplementary Figure 2. Connectedness between mOTUs based on the presence (1, green) or absence (0, gray) of CRISPR spacer clusters targeting the same vOTUs and PTUs between them.*

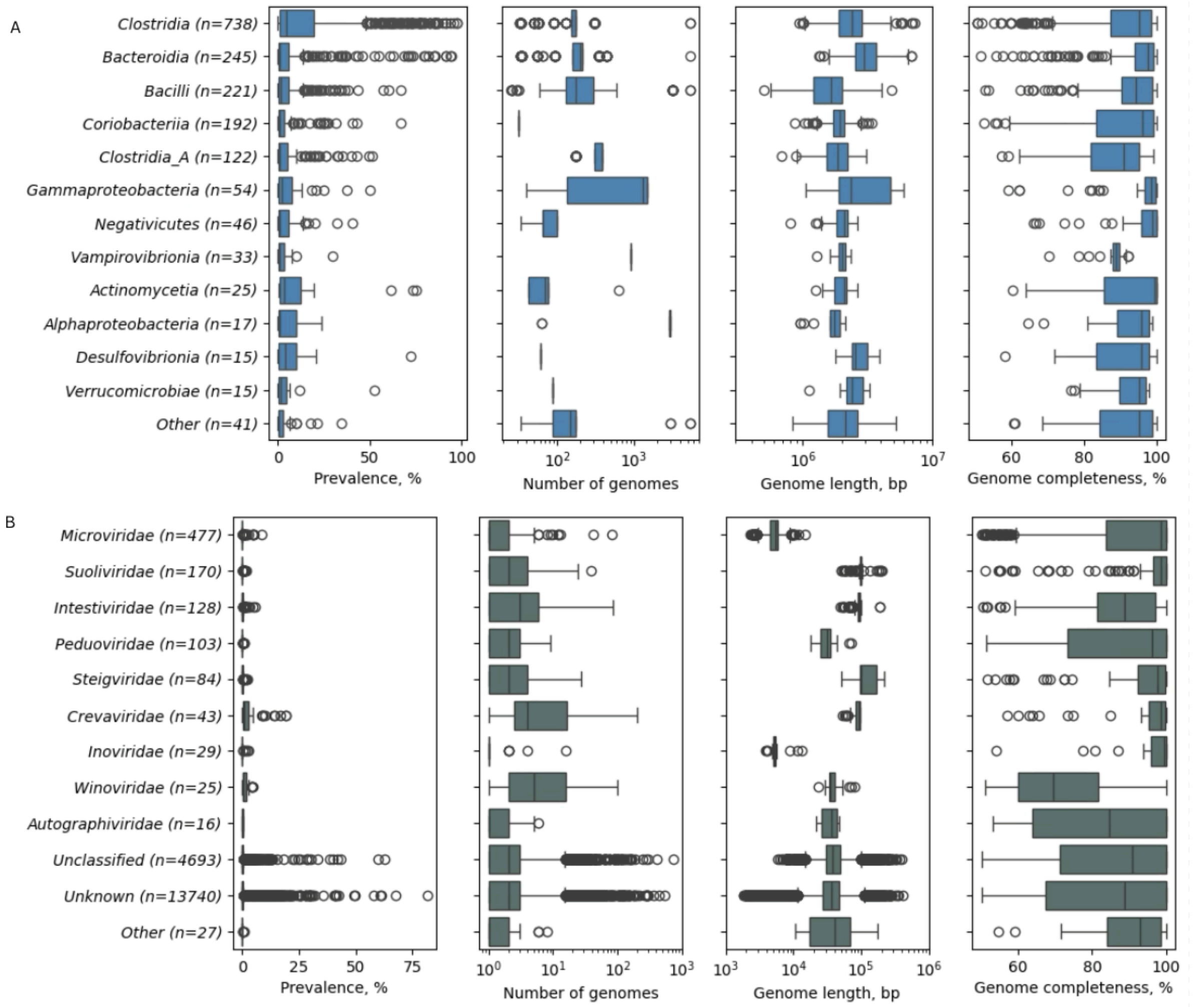


*Supplementary Figure 3. Bacteriome/archaeome and virome overview. A) Prevalence, genome length and completeness of mOTUs. B) Prevalence, genome length and completeness of vOTUs*

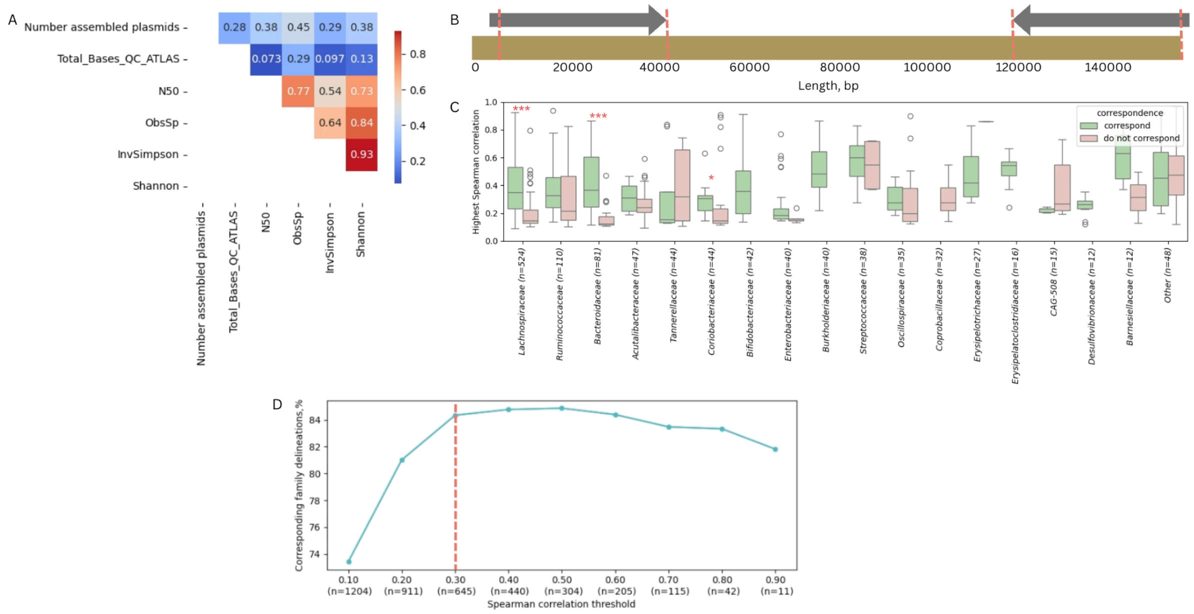


*Supplementary Figure 4. Technical characteristics of the plasmidome data A) Pearson's correlation between number of assembled plasmids, sequencing depth (Total bases QC), assembly quality (N50), and bacterial diversity (Observed species, Inversed Simpson and Shannon indices) B). Plasmid assembly circularity issue. Homology regions between CRCbiome-PTU_02517 (Unknown host family) and its best matching IMG/PR reference plasmid sequence. Brown - CRCbiome-PTU_02517 complete sequence, Red dashed lines - homology regions (99.3% identity), grey - IMGPR_plasmid_3300029875_000076 complete sequence, arrow depicts 3'-end. C) Spearman correlation coefficient between PTU and mOTU stratified by whether PTU host prediction and mOTU taxonomy delineation correspond (green) or do not correspond (grey) D). Fraction of PTUs with predicted host family corresponding to that of highest-correlating mOTU given the minimal allowed Spearman correlation value.*

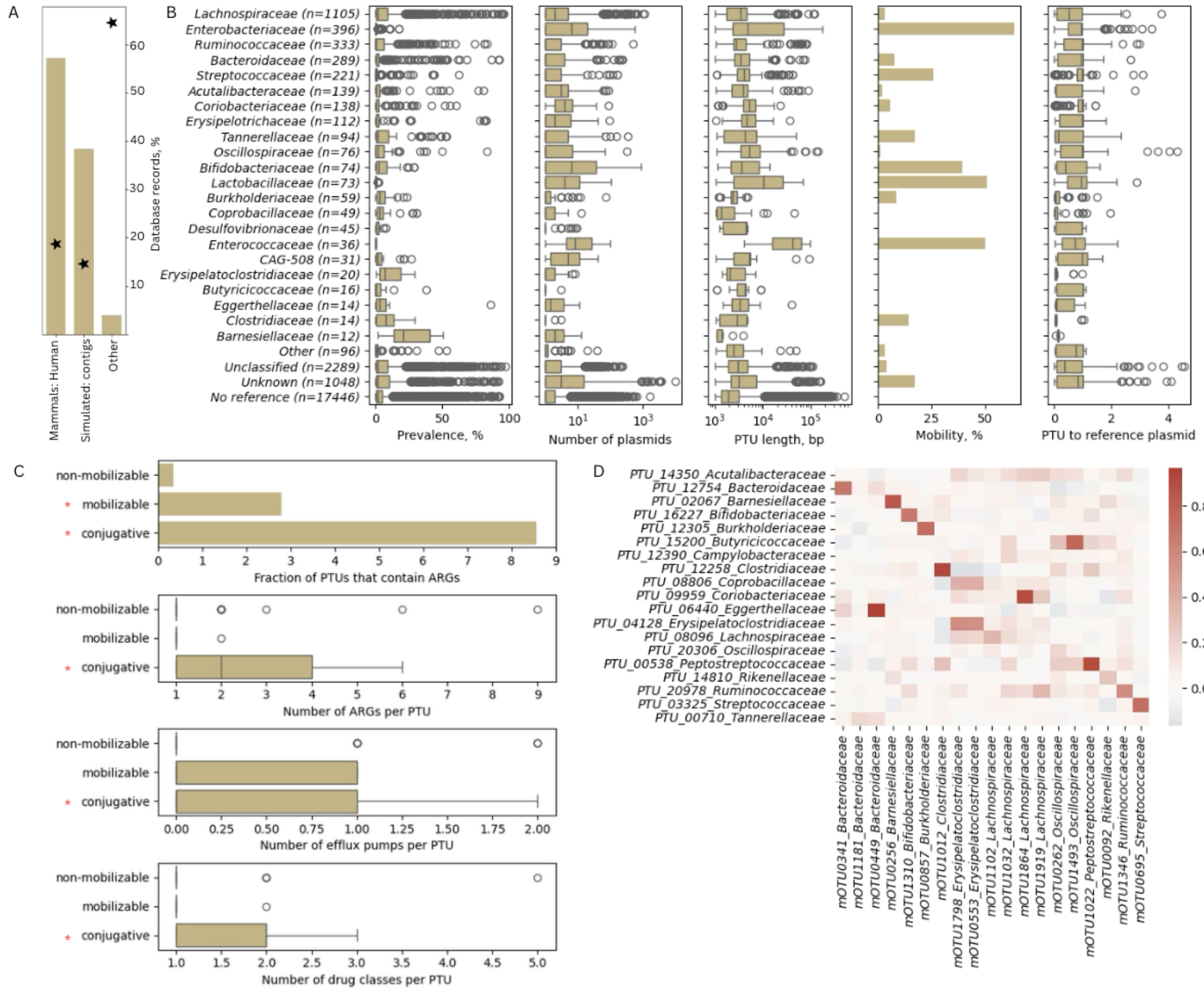


*Supplementary Figure 5. Plasmidome overview. A) Source types of IMG/PR reference plasmid sequences that had the closest homology to CRCbiome PTUs. Bars - only plasmids matching to PTUs from the CRCbiome dataset, stars - complete IMG/PR database; B) General characteristics of the CRCbiome plasmidome dataset stratified by taxonomy of the predicted bacterial host, ordered most to least frequent family delineation. No reference: No matching reference plasmid in IMG/PR database; Unclassified: IMG/PR reference lacks predicted host taxonomy assignment; Unknown: host taxonomy is inferred to a higher taxonomic level than family. Panels (left to right): Prevalence of PTUs; Number of plasmids in PTUs within a family; PTU length; Percent of mobile (mobilizable and conjugative) PTUs in each family; Ratio between PTU length and best matching IMG/PR reference plasmid length; C) Mobility and antibiotic resistance genes load of PTUs. *p<0.05 D) Correspondence between PTU host delineation and mOTU taxonomy based on Spearman correlation between PTUs and strongest correlated mOTUs. For visualization purposes, only the PTU with the highest prevalence within the PTU predicted host family is depicted. *padj < 0.001*

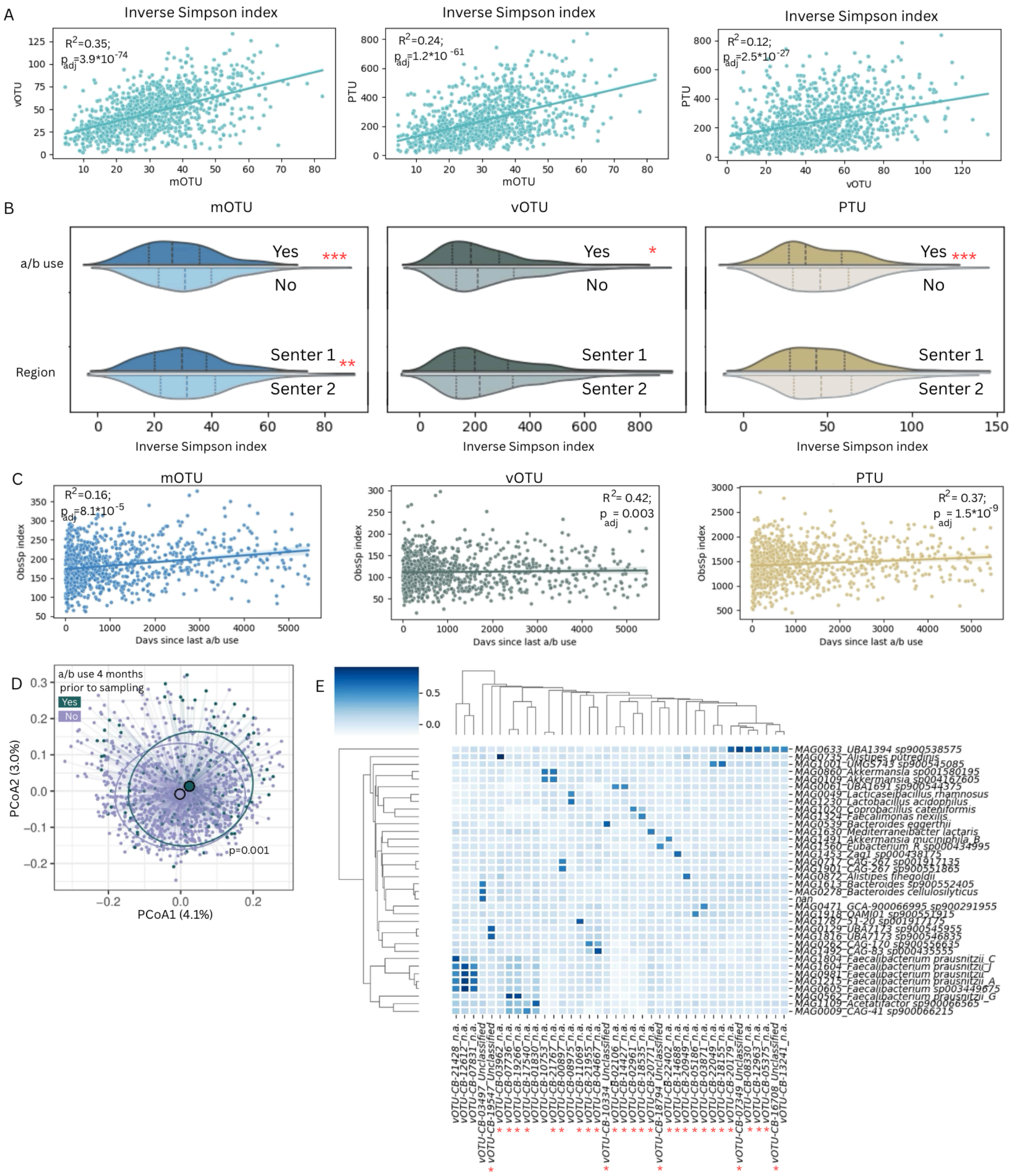

*Supplementary Figure 6. Extended microbiome characteristics A) Correlation between Inversed Simpson index of mOTUs, vOTUs and PTUs. B) Difference in Inversed Simpson index with regard to prescribed antibiotics use within 4 months prior to sampling (top) and residence region (bottom) for mOTUs (blue), vOTUs (green) and PTUs (blown). C) Linear regression between microbial diversity and days passed since last antibiotics prescription for mOTUs (blue), vOTUs (green) and PTUs (blown). The model is adjusted for the sequencing depth. D). Principal Coordinate Analysis based on extended microbiome Bray-Curtis index (mOTUs, vOTUs and PTUs together) stratified by antibiotics use within 4 months prior to sampling. E) Heatmap showing spearman correlation between relative abundance of mOTUs and vOTUs. For visualization purposes, only mOTUs/vOTUs with any |rho|>0.5 are included. Red asterisks mark vOTUs that include viral genomes incorporated into MAG contigs. The cluster of Faecalibacterium mOTUs with varying viral correlation profiles is highlighted.*
